# Rapid and continuing regional decline of butterflies in eastern Denmark 1993-2019

**DOI:** 10.1101/2023.04.03.535267

**Authors:** Emil Blicher Bjerregård, Lars Baastrup-Spohr, Bo Markussen, Hans Henrik Bruun

**Affiliations:** Department of Biology, University of Copenhagen, Universitetsparken 15, Copenhagen, Denmark; Department of Mathematical Sciences, University of Copenhagen, Universitetsparken 5, Copenhagen, Denmark

**Keywords:** Butterfly monitoring, extinction debt, grassland, habitat loss, Hesperiidae, Lycaenidae, Nymphalidae, Papilionidae, Pieridae, peat bog, woodland

## Abstract

1. Many butterfly populations respond negatively to land-use intensification in human-dominated landscapes. However, networks of protected sites have been established with the aim to halt species loss.
2. We undertook annual surveys of all populations of 22 uncommon butterfly species in eastern Denmark during the period 2014-2019 and compared to a systematic atlas survey done 1989-1993, in order to assess trends in regional occupancy of species.
3. Three out of 22 species went regionally extinct between 1993 and 2015. One species sustained a single population through the study period. Logistic regression for the remaining 18 species showed 10 to be in strong decline from 1993 to 2015, two showed a declining trend and six had stable trends. For all species except one, the declining trend continued 2015-2019. For five species, a sustained strong decline was evident.
4. In 1993, the total count of populations for all 22 butterfly species was 565, which by 2019 had declined to 158 populations (a 72 % loss over 26 years). From 2015 to 2019, the total count of populations further shrank from 200 to 158 (a 21 % decline over just four years).
5. Legal protection of areas (Natura 2000 and Danish Nature Protection Act §3) was, unexpectedly, not associated with lower probability of local extinction for butterfly population. The observed sustained decline across species suggests an overall low efficiency of the network of protected sites, probably due to a combination of misguided management regimes and payment of extinction debts from the past.

## Introduction

Habitat change caused by human land-use is well documented as the main driver of biodiversity change worldwide (IPBES 2019; Newbold et al. 2015). Anthropogenic impact on natural habitats has taken place through millennia (Ellis et al. 2021), but the magnitude and intensity have dramatically risen during the industrial era (Mihoub et al. 2017). Insects are among the organisms suspected to suffer from widespread declines (van Klink et al. 2020; Wagner 2020).

Butterflies have been shown to be particularly sensitive to anthropogenic land-use intensification and the majority of species are thought to decline faster than other insect groups (Forister et al. 2021; Warren et al. 2021; Wepprich et al. 2019). Declining trends for many species of butterfly have been observed across northern Europe. In the Netherlands, a 84 % long-term decline (1890-2017) for 71 butterfly species was estimated from annual transect data, with a 50 % decline in recent years (post 1992) alone (van Strien et al. 2019). Similarly in the UK, specialist butterfly species continue to show declining trends, i.e. 7.7 % and 13.9 % decline for the periods 1994-2001 and 2002-2009, respectively (Brereton et al. 2011). Similar patterns of general decline have been observed over larger parts of Europe (van Swaay et al. 2019; Warren et al. 2021).

Since 1990, Butterfly Conservation Europe has run The European Butterfly Monitoring Scheme (eBMS), but Denmark has not been covered by this citizen science effort. On this background, (Eskildsen et al. 2015) compiled a vast data set on Danish butterfly diversity 1900-2012. They found strong declines, including regional extinction of 10 % of the entire butterfly fauna, in particular species associated with open woodlands and forest glades. Species that are sedentary habitat specialists were found to decline disproportionately strongly, while mobile generalists remained about stable in regional occupancy. Their results indicated that in Denmark, butterfly habitats associated with the forest-grassland ecotone and with open woodland declined strongly and relatively early in the 20^th^ Century, whereas open grasslands and heathlands generally deteriorated in the second half of the 20th Century.

Denmark has seen major land-use changes over the past century, which has driven changes in communities of plants, animals and fungi. The most significant land-use changes were: 1) drainage of moist and wet biotopes, 2) active afforestation and passive scrub overgrowth due to ceased grazing, and 3) landscape-wide increased availability of plant nutrients (Finderup Nielsen et al. 2021). The total area under rotational farming has decreased during the last 50 years (Levin and Normander 2008), so most recent land-use changes may be attributed to intensified plantation forestry and abandonment of livestock grazing in the landscape at large. From a butterfly perspective, many key habitat types have seen continuous decline in both area and habitat quality.

Protected areas (PA) have been designated under Danish legislation since 1917. In 1992, a revised nature conservation act stipulated status quo of all remaining areas of grassland, heathland, mire, salt marsh, fresh meadow and other open habitat types. The Habitats Directive of the European Union was later implemented and stipulated protection of certain types of forested habitats as well. One could expect the decline in occupancy of many butterfly species demonstrated by Eskildsen et al. (2015) to have stabilized after the implementation of strengthened nature conservation acts. The efficiency of protected areas for biodiversity conservation is a topic of both scientific and practical interest (e.g. Geldmann et al. 2013; Rada et al. 2019).

We aimed at assessing the occupancy trends in recent decades (1993-2019) for butterflies in eastern Denmark, excluding highly mobile and generalist species, i.e. focussing on 22 less common habitat-specialist species, all having restricted distribution ranges and being sufficiently sedentary to form distinct populations enabling use of presence-absence data. Specifically, we aimed at:

1. assessing if the declining trend for Danish butterfly populations continued in the time period 1993-2015 and if any trend would continue during 2015-2019; 2) assessing trends for groups of species associated with broadly defined habitat types in order to relate trends to land-use change and, 3) assessing to which degree butterfly populations had lower probability of decline in legally protected areas.

## Materials & Methods

The study area was eastern Denmark, i.e. the main islands of Zealand, Lolland, Falster and Møn with smaller coastal islands, i.e. a total land area of c. 9000 km^2^, dominated by arable farmland and with appreciable urban areas and managed forests (Levin et al. 2017). We focussed on sites occupied by populations of the following 22 butterfly species: *Carterocephalus silvicola* (Meigen, 1829), *Erynnis tages* (L., 1758), *Pyrgus malvae* (L., 17858), *Pyrgus armoricanus* (Oberthür, 1910), *Hesperia comma* (L., 1758), *Lycaena hippothoe* (L., 1761), *Callophrys rubi* (L., 1758), *Phengaris arion* (L., 1758), *Plebejus argus* (L., 1758), *Plebejus idas* (L., 1761), *Aricia artaxerxes* (Fabricius, 1793), *Agriades optilete* (Knoch, 1781), *Cyaniris semiargus* (Rottemburg, 1775), *Polyommatus amandus* (Schneider, 1792), *Brenthis ino* (Rottemburg, 1775), *Speyeria aglaja* (L., 1758), *Fabriciana adippe* (Denis & Schiffermüller, 1775), *Fabriciana niobe* (L., 1761), *Boloria aquilonaris* (Stichel, 1908), *Boloria selene* (Denis & Schiffermüller, 1775), *Boloria euphrosyne* (L., 1758) and *Coenonympha tullia* (Müller, 1764). This selection excluded mobile and habitat-generalist species. A few casual or climate-driven recent immigrant species, which have had relatively short-lived presences in the study area, viz. *Lycaena tityrus* (Poda, 1761) and *Heteropterus morpheus* (Pallas, 1771), were also excluded.

Approximately 16 % of Denmark’s land area enjoys some form of legal nature protection, with Clause 3 (§3) of the Danish Nature Protection Act (effective as of 1993) and Natura 2000 (EC Habitats Directive 1992, implemented legally and administratively in Denmark in a protracted process c. 2000-2010) being the most important. Clause 3, which prohibits land-use intensification in non-forest habitats, viz. lakes, salt meadows, fresh meadows, bogs, mires, heathlands and dry grasslands, covers 10.5 % of the land area including lakes. The Natura 2000 network covers 9 % and includes open and forested habitat types. The two schemes together, excluding overlap in areas covered by both, sum to 16.1 % of the land area (Biodiversitetsrådet 2022).

### Atlas of Danish Butterflies 1989-1993

During a five-year period from 1989-1993, a citizen science project was conducted with volunteers recording butterfly populations across Denmark. This was effectively the first atlas survey of Danish butterfly species (Stoltze 1996). For the study area and species considered here, the atlas data consist of 2146 records of 25 species at 364 sites. The recording activity was distributed over all four years in order to obtain the best possible coverage of the whole country, so sites were not necessarily visited each year. Thus, for the present analysis, the atlas data are considered one snapshot of species occurrence. We assume no population trend during the time period and thus assume all populations to have been present by 1993. For the study area and species considered here, the data consist of 565 presence records of 22 species at 214 sites in a single time slice.

### Field survey 2014-2019

During the six-year period 2014-2019, Emil Blicher Bjerregård and Magnus Vest Hebsgaard aimed at making an annual occupancy status of uncommon butterfly species on Zealand, Lolland, Falster and Møn by 1) visiting all known populations at least once a year during the flight periods of all target species, 2) strategically searching for overlooked populations at apparently suitable sites and sites having harboured populations prior to 1989. The aim was to obtain exhaustive presence-absence data for the selected 22 extant species within the study area. It would not have been feasible to apply this methodology to recording of widely distributed species. Common to most of the studied butterfly species is a high degree of visibility in their adult stage. This makes it relatively time-economical to ascertain the presence or absence of a butterfly species, whereas attempts to estimate population sizes would have multiplied the necessary time consumption. Although a species at a given site was not found to be present in a given year, it would still be the target of searches the following years, in order to increase data credibility. In planning annual field surveys, small populations were given the highest priority on days of benign weather conditions right at the peak of the target species’ flight time in order to increase the probability of recording presence. Conversely, large populations, the presence of which is more readily detected even at the beginning or end of the target species’ flight time, were fitted into the schedule or visited during days of less favourable weather conditions. For the study area and species considered here, the data consist of 525 site × year presence records of 19 species at 214 sites (three species had gone regionally extinct after 1993).

## Data preparation

A number of specific records in the atlas data were corrected, based on knowledge obtained later than 1993. Most importantly, 64 absence records were changed to presence in specific years, if the focal species at the given site was recorded in later years, effectively assuming that local extinction and subsequent re-immigration was considered much less likely than oversight, e.g. due to the lack of neighbouring populations within a reasonable distance. Also, 121 records were deleted using ancillary data from Bugbase (Danish Lepidopterological Society). Four records were omitted as they were considered cases of a stray individuals not representing a population (Table S3; Supplementary Materials).

Each species was assigned to one broadly defined habitat category, i.e. inland grassland, coastal grassland, open woodland and peat bog. The ‘inland grassland’ species group comprised three typical grassland species with known distribution ranges mainly in the inland. The ‘coastal grassland’ group contained five species, which in eastern Denmark are confined to the coastal zone. The ‘open woodland’ and ‘peat bog’ groups are both well circumscribed in the region (Supplementary Materials, Table S1). In the analysis of site protection, we considered the species groups inland grassland, coastal grassland and peat bog as an ‘open habitats’ group, for which Clause 3 (§3) of the Danish Nature Protection Act is relevant.

All data from the 1989-1993 atlas survey were assigned to a time slice in the year 1993. Data from the 2014-2019 survey were partitioned into three separate time slices, viz. 2014-2015 (“2015” henceforth), 2016-2017 (“2017”) and 2018-2019 (“2019”). This was done in order to ensure data reliability and remove any effect of potential single-year oversights of population presences.

### Assignment of site identity to old records

Prior to analysis, site names in the atlas data and the recent survey were standardized. First, sites in the sense used in the present study were delineated. A site was defined as a contiguous area with only one of the mentioned habitat types. Neighbouring areas with different habitat types were assigned to separate sites. The delineation was done at a scale sufficiently coarse to ensure that atlas data, which lacked exact geo-referencing, were matched irrespectively of the toponym used by the recorder. In almost all cases, there was no doubt about the exact geographic origin of a record, because of the fragmented character of eastern Danish landscapes with natural areas separated in a matrix of arable land, and because most butterfly populations have been known to collectors for generations. However, the coarse scale allowed the crucial inclusion of records only assigned the name of a forest or similar. In the case of peat bogs embedded in forest, several neighbouring bogs were all assigned the name of the forest. The same was the case for forest meadows. Site standardization resulted in a total of 214 sites included for analysis. Examples of site delineation are given in Fig. S1 and S2 (Supplementary Materials). All changes from the original site names are documented in Table S2 (Supplementary Materials).

## Statistical analysis

The species *A. artaxerxes, H. comma* and *F. niobe* were omitted from statistical analysis, because they were only observed at a single site each in 1993 and went extinct in the study area prior to 2015. Similarly, *C. tullia* was omitted at this stage, because it was observed at a single site only throughout the sampling years. For the remaining 18 species, we performed four collections of statistical analyses separately for each species. Finally, the resulting 18 analyses were combined across species in order to provide overall conclusions and to correct for the associated multiple testing problem. Below, we will give the details of the analytical steps.

The first collection of statistical analyses was a logistic regression for *Y* (1 = presence, 0 = absence) of a butterfly species across the 214 sites in the four time slices in the period 1993 - 2019. We had 4 * 214 = 856 observations of *Y* and the predictor variables Year and Site, and we modelled the odds for presence at a site via

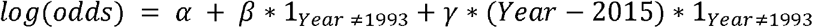

Here, the parameter *α* quantifies the *log(odds)* for presence among the sites in 1993, *β* quantifies the *log(odds ratio)* between year 2015 and year 1993, and *γ* quantifies the *log(odds ratio)* per calendar year since 2015. Moreover, we corrected for potential dependence between the four observations within the same *Site* using the *Generalized Estimation Equation* methodology (Liang and Zeger 1986) via the *geepack* package version 1.3.4 (Højsgaard et al. 2006) in the statistical software *R* version 4.2.0 (R Core Team 2022). The regression model stated above entails an assumption of constant odds ratios per calendar year from 2015 to 2019. This assumption was tested by a Goodness-of-Fit test against the saturated model:

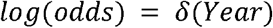

Upon validation of the regression models, we provide estimates of the odds ratios for presence between 2015 and 1993 and per calendar year from 2015 to 2019. Moreover, we provide hypothesis tests for the null hypothesis of odds ratios being equal to 1, i.e. corresponding to no change in the odds for presence.

The three other collections of statistical analyses were concerned with the potential effect of legal nature protection: 1) *Clause 3* (§3) of the Danish Nature Protection Act; 2) *Natura 2000*; 3) The combination of the two, i.e. *Natura 2000* status for occurrence sites of *open woodland* species and *Natura 2000* and/or *Clause 3* for occurrence sites of the remaining species, which will be called Protected henceforth. One site in the 1993 atlas data (named after the town Rønnede) could not be relocated with certainty and had to be omitted from this analysis. For these collections of statistical analyses, we performed two-sided Fisher’s Exact test and estimated odds ratios on the following three cross tabulations:

1. *Presence/absence* in 1993 vs. protection status *Yes/No* among all 213 sites.
2. Protection status *Yes/No* vs. *Presence/Absence* in 2015 among the sites with the butterfly species present in 1993.
3. Protection status *Yes/No* vs. *Presence/Absence* in 2019 among the sites with the butterfly species present in 2015.

Since both *Clause 3* and *Natura 2000* were implemented between 1993 and 2015, cross tabulation 1 mainly quantifies the odds ratio for selection of sites for legal nature protection, whereas the odds ratios in cross tabulations 2 and 3 quantify the subsequent effect of legal protection on extinction probability of butterfly populations. For cross tabulations 2 and 3, the two-sided test makes no a *priori* assumption that legal protection cannot possibly have negative effects on population survival.

The results for the separate analyses done per species are collected in combined tables with estimates and 95 pct confidence intervals for the odds ratios and p-values for the null hypothesis of no change (i.e. odds ratio equal to 1). Results are written in italics for p-values < 0.05 and in boldface for Holm adjusted p-values < 0.05. The Holm (1979) adjustment ensures Type I control of the Family Wise Error Rate. Thus, it provides a statistical analysis across the 18 butterfly species. In addition, we combined p-values across the investigated butterfly species in more ways:

1. We performed the Too Many Too Improbable test (Mogensen and Markussen 2022) for the joint null hypothesis of no effect for all butterfly species.
2. We performed the Closed Testing Procedure (Marcus et al. 1976) in order to obtain a 95 pct confidence set for the number of false null hypotheses (Goeman and Solari 2011) among the species with uncorrected p-values < 0.05. This is done based on the TMTI test using the TMTI package version 0.1.0 in R (Mogensen 2021).

These combined analyses were done both for all investigated species, and separately within the four habitat-defined species groups, viz. coastal grassland, inland grassland, peat bog and open woodland. The latter was done to investigate whether trends differed between habitat types.

## Results

During the quarter-of-a-century period from the Atlas of Danish Butterflies in 1993 to the end of our survey in 2019, our data demonstrate an overall decline by 72 % in the total number of populations of the focal 22 butterfly species (from 565 to 158 populations in total). The decline for the same 22 species during the four-year period 2015-2019 was 21 % (from 200 to 158 populations).

For all species, the odds ratio for species’ occurrence in 2015 compared to 1993 was estimated to be below 1, which means that all species have declined (Table 1). Under control of the family-wise error rate, the declining trend was statistically significant at the 5 pct level for ten species out of 18, viz. *adippe, aglaja, amandus, argus, euphrosyne, hippothoe, idas, ino, malvae, selene*. For further two species, *optilete* and *semiargus*, individual tests of decline were significant at the 5 pct level. Among these 12 species, the closed testing procedure provided 95 pct confidence that at least 11 species showed declines from 1993 to 2015.

**Table 1.** Results of logistic regressions across 18 butterfly species. GoF column provides Goodness-of-Fit tests for the assumption of constant odds ratios per year from 2015 to 2019. Estimates and 95 pct confidence intervals (in the parenthesis) for the odds ratios, and two-sided p-values for the null hypothesis of odds ratio equal to 1, are adjusted for correlation within sites via GEE. Italics or boldface annotations imply significance on the 5 pct level within species, and boldface imply significance on 5 pct level across species (Holm adjustment). Both are given with asterisks as visual cues. If a species was present only at a single site in 2015 and later, then neither odds ratio nor p-value are stated for the trend between 2015 and 2019.

| Species | GoF | Odds ratio (2015 vs. 1993) | p-value | Odds ratio (2015-2019) | p-value |
| --- | --- | --- | --- | --- | --- |
| *adippe* | 0.2298 | **0.23 (0.14 ; 0.39)** | **0.0001\*** | 0.88 (0.79 ; 0.99) | 0.0241* |
| *aglaja* | 0.0819 | **0.23 (0.13 ; 0.38)** | **0.0001\*** | 0.94 (0.87 ; 1.02) | 0.0922 |
| *amandus* | 0.0817 | **0.14 (0.08 ; 0.22)** | **0.0001\*** | 0.95 (0.89 ; 1.01) | 0.0906 |
| *aquilonaris* | 0.7265 | 0.77 (0.42 ; 1.39) | 0.3834 | 0.77 (0.54 ; 1.09) | 0.1379 |
| *argus* | 0.5477 | **0.57 (0.40 ; 0.79)** | **0.0009\*** | 0.98 (0.92 ; 1.05) | 0.5647 |
| *arion* | 1.0000 | 0.50 (0.12 ; 2.01) | 0.3260 | NA | NA |
| *armoricanus* | 0.9089 | 0.83 (0.57 ; 1.22) | 0.3428 | 1.09 (0.96 ; 1.23) | 0.1543 |
| *euphrosyne* | 0.3180 | **0.13 (0.05 ; 0.34)** | **0.0001\*** | 0.93 (0.79 ; 1.09) | 0.3386 |
| *hippotoe* | 0.8124 | **0.36 (0.23 ; 0.55)** | **0.0001\*** | 0.90 (0.81 ; 0.99) | 0.0252* |
| *idas* | 0.2808 | **0.63 (0.47 ; 0.86)** | **0.0024\*** | 0.94 (0.88 ; 1.00) | 0.0396* |
| *ino* | 0.5910 | **0.31 (0.20 ; 0.49)** | **0.0001\*** | 0.87 (0.77 ; 0.98) | 0.0153* |
| *malvae* | 0.6090 | **0.24 (0.16 ; 0.38)** | **0.0001\*** | 0.95 (0.89 ; 1.01) | 0.0840 |
| *optilete* | 0.1601 | 0.41 (0.21 ; 0.81) | 0.0097* | 0.87 (0.70 ; 1.08) | 0.1913 |
| *rubi* | 0.1556 | 0.86 (0.72 ; 1.02) | 0.0691 | 0.98 (0.94 ; 1.01) | 0.1503 |
| *selene* | 0.1138 | **0.15 (0.09 ; 0.23)** | **0.0000\*** | 0.93 (0.87 ; 1.00) | 0.0271* |
| *semiargus* | 1.0000 | 0.07 (0.00 ; 0.52) | 0.0097* | NA | NA |
| *silvicola* | 1.0000 | 0.33 (0.06 ; 1.65) | 0.1768 | NA | NA |
| *tages* | 1.0000 | 0.50 (0.12 ; 2.01) | 0.3260 | NA | NA |

For the period 2015 - 2019, all but one species had estimated odds ratios below 1, indicating overall decline. The exception was *armoricanus*, which had an estimated (but not significant, p = 0.154) odds ratio above 1. However, none of the changes were significant under control of the family wise error rate. The TMTI test, on the other hand, provided strong evidence that at least one species was declining (p < 0.0001), and 95 pct confidence that at least three of the species *adippe, hippothoe, idas, ino, selene* had declined. All species met the assumption of constant odds ratio per calendar year from 2015 to 2019 as validated by the Goodness-of-Fit test (Table 1).

The declining trend in occupancy was pertinent to all four habitat-defined groups of species, both 1993-2015 and 2015-2019 (Fig. 1; Table 2) and with comparable variation among species in all four groups.

**Fig. 1:**
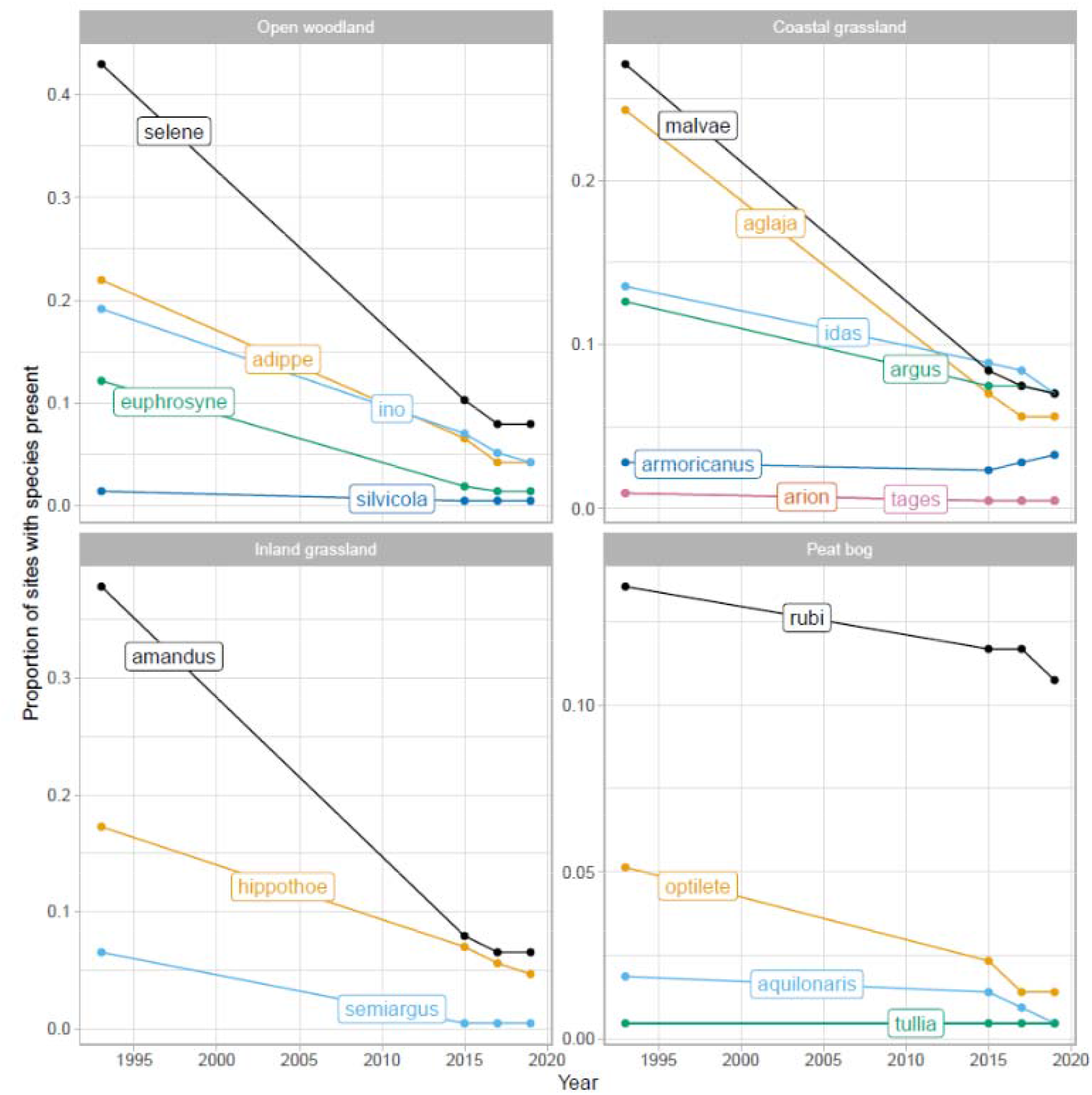
Trends in occupancy of 19 uncommon butterfly species in Eastern Denmark 1993-2019, with species allocated to broadly defined habitat categories. Please note that the ordinate axes have different units in order to maximize visual resolution (for a version of the figure with identical units – see Fig. S5). Coenonympha tullia, which occurred at a single site throughout the sampling period, is shown in the figure, but was omitted from statistical analyses. The species artaxerxes, comma and niobe were observed at a single site each in 1993, but went extinct in the study area prior to 2015 and are not shown.

**Table 2.** Occupancy change per species group defined by habitat affinity (group size in brackets). For each of the two time periods, the p-value for the null-hypothesis of no occupancy change is given. Similarly, the lower confidence limit for the number of species with a decline among those with a marginal p-value < 0.05 of occupancy change is given.

| Habitat-defined species group | 1993-2015 |  | 2015-2019 |  |
| --- | --- | --- | --- | --- |
|  | p-value | Lower CI | p-value | Lower CI |
| Open woodland (5 spp) | < 0.0001 | 4 | < 0.001 | 2 |
| Coastal grassland (7 spp) | < 0.0001 | 4 | 0.011 | 0 |
| Inland grassland (3 spp) | < 0.0001 | 3 | 0.016 | 1 |
| Peat bog (3 spp) | 0.036 | 1 | 0.019 | 0 |
| All 18 species | < 0.0001 | 11 | < 0.0001 | 3 |

For most of the focal butterfly species, 1993 occurrence sites had higher probability of later being selected for legal protection than did vacant sites (Table 3). This was the case for the 13 species associated with non-forest habitats in relation to the Danish Clause 3 protection of open habitat types (Table S4, Supplementary materials), as well as for all species in relation to Natura 2000 (Table S5, Supplementary materials) and for the combination of the two protection schemes (Table 3). In contrast, the evidence that legal protection by Clause 3 of the Danish Nature Protection Act and Natura 2000 resulted in lower extinction rate within protected areas than outside was very weak at best. We did not find evidence for efficiency of Clause 3 in any time slice comparison, i.e. 1993-2015 (joint test: p=0.35) and 2015-2019 (joint test: p=0.96). Similarly for Natura 2000 and for both periods, i.e. 1993-2015 (joint test: p=0.68) and 2015-2019 (joint test: p=0.99), and for the combination of Clause 3 and Natura 2000, 1993-2015 (joint test: p=0.10) and 2015-2019 (joint test: p=0.99).

**Table 3.** Fisher’s Exact test investigating protected area status (Clause 3 and Natura 2000 combined) across 13 butterfly species associated with open habitats. All tests are two-sided. Italics or boldface imply significance at the 5 pct level within species, and boldface implies significance at the 5 pct level across species. Both are given with asterisks as visual cues. If all sites with the species present from the starting year had the same legal protection status, then neither odds ratio nor p-value are given.

| Species | **Odds ratio**<br>**(Yes vs No)** | **p-value** | **Odds ratio**<br>**(2015 vs 1993)** | **p-value** | **Odds ratio**<br>**(2019 vs 2015)** | **p-value** |
| --- | --- | --- | --- | --- | --- | --- |
| aglaja | 1.82<br>(0.89 ; 3.71) | 0.0796 | *4.51 (0.98 ; 24.67)* | *0.0401\** | 1 (0.01 ; 25.35) | 1.0000 |
| amandus | **2.8 (1.46 ; 5.43)** | **0.0011\*** | 3.22 (0.94 ; 12.09) | 0.0516 | 4.49 (0.19 ; 322.88) | 0.5148 |
| aquilonaris | 7.72<br>(0.61 ; 411.64) | 0.0719 | 0 (0 ; 116.79) | 1.0000 | Inf (0.01 ; Inf) | 1.0000 |
| argus | **4.52 (1.82 ; 11.63)** | **0.0005\*** | 0.74 (0.11 ; 4.53) | 1.0000 | Inf (0.25 ; Inf) | 0.1751 |
| arion | Inf (0.47 ; Inf) | 0.0811 | 2 sites with presence in 1993 all protected | NA | 1 sites with presence in 2015 all protected | NA |
| armoricanus | 13.3<br>(1.44 ; 639.48) | *0.0080\** | 0 (0 ; 194.41) | 1.0000 | 0 (0 ; Inf) | 1.0000 |

| Species | Odds ratio<br>(Yes vs No) | p-value | Odds ratio<br>(2015 vs 1993) | p-value | Odds ratio<br>(2019 vs 2015) | p-value |
| --- | --- | --- | --- | --- | --- | --- |
| hippotoe | 1.67<br>(0.73 ;<br>3.71) | 0.2292 | 0.73 (0.14 ;<br>3.42) | 0.7380 | 2.51 (0.15 ;<br>163.45) | 0.6004 |
| idas | **9.32**<br>**(3.64 ;**<br>**26.27)** | **0.0001\*** | 2.42 (0.33 ;<br>18.16) | 0.3905 | 1.31 (0.02 ;<br>25.74) | 1.0000 |
| malvae | **3.08**<br>**(1.55 ;**<br>**6.17)** | **0.0007\*** | 3.26 (0.9 ;<br>13.02) | 0.0502 | 4.93 (0.2 ;<br>352.16) | 0.2451 |
| optilete | 4.75<br>(1.16 ;<br>23.05) | 0.0143* | 3.52 (0.17 ;<br>260.91) | 0.5455 | Inf (0.04 ; Inf) | 0.4000 |
| rubi | **9.32**<br>**(3.64 ;**<br>**26.27)** | **0.0001\*** | 0.86 (0.01 ;<br>13.05) | 1.0000 | 0 (0 ; 14.12) | 1.0000 |
| semiargus | 1.42<br>(0.36 ;<br>4.95) | 0.5491 | 0 (0 ; Inf) | 1.0000 | 1 sites with<br>presence in<br>2015 all un-<br>protected | NA |
| tages | Inf (0.47 ;<br>Inf) | 0.0811 | 2 sites with<br>presence in<br>1993 all<br>protected | NA | 1 sites with<br>presence in<br>2015 all<br>protected | NA |

Our data demonstrate plummeting occupancy of nearly all uncommon butterfly species in eastern Denmark in recent decades. Further, the data show that – despite relevant sites being selected for legal nature protection – neither national nor EU legislation have been efficient enough to prevent local extinction of uncommon butterflies, even from protected areas. Thus, despite formal protection being in place for decades, the decline of butterflies continues unabated.

The species studied were all relatively sedentary species with rather clear habitat affiliations. Our analysis does not say anything about mobile habitat generalists but, apparently, these species seems not to decline systematically in the study area. By necessity, we also excluded the 11 species known from the study area during the 20^th^ Century, but now all extinct from Denmark at large (*Parnassius mnemosyne, Leptidea sinapis, Leptidea juvernica, Hamearis lucina, Satyrium pruni, Satyrium ilicis, Lycaena dispar, Limenitis populi, Melitaea diamina, Coenonympha arcania and Coenonympha hero*) and the three species that went extinct in eastern Denmark after our starting point in 1993, but which maintain populations elsewhere in Denmark (*Hesperia comma, Aricia artaxerxes and Fabriciana niobe*). Thereby, we see our analysis as being balanced between conservative and liberal. The overall trend is one typical of biotic homogenization, which means that regionally uncommon species decline or go extinct, while already widespread species increase or at least remain about constant (Finderup Nielsen et al. 2019; McKinney and Lockwood 1999), although the presented analysis did not deal with that question directly. Inclusion of widespread species or focus on local richness seems unlikely contribute more clarity (Larsen et al. 2018).

The observed trend is similar to long-term trends from similar countries, such as the UK, where 80 % of butterfly species have decreased in abundance or distribution, or both, since the 1970s, in particular habitat specialist like the suite of species studied here (Fox et al. 2023). The sustained decline of many of the focal species in eastern Denmark, even in the last decade, appears to be more dramatic than elsewhere in Europe (Habel et al. 2022a; van Swaay et al. 2019; Warren et al. 2021).

The observed trend was not restricted to one particular habitat type, but prevailed in grasslands, open woodlands and peat bogs, each with their set of more or less narrow habitat specialists. Five of our focal species, as well as several of the butterfly species now extinct in Denmark, are affiliated with open woodlands and forest meadows (Habel et al. 2022b), which to a large extent have been lost to the intensification of plantation forestry and to less extent to game management. Similarly, drainage as part of intensified forestry has negatively affected peat bogs, which in eastern Denmark mainly occur in forest landscapes. Coastal and inland grasslands in eastern Denmark have mainly changed due to summer grazing at high stocking densities or, even more widespread, abandonment of grazing altogether (Finderup Nielsen et al. 2021). Some sites have been lost to urbanization, despite the general protection of grasslands implemented by Clause 3 of the Danish Nature Protection Act (examples shown in Fig. S3 and S4, Supplementary materials).

The 1993 presence of one or more populations of uncommon butterfly species was a strong predictor of sites being selected for legal protection under either or both the Danish Nature Protection Act and the EU Habitats Directive. Although this observation does not necessarily imply that butterfly occurrences were important parts of the basis for decisions, it does imply that conservation officers were well aware of extant biodiversity values in general terms. Only one of the included butterfly species, *Maculinea arion*, is listed on Annex 4 of the EU Habitats Directive as a species of community interest in need of strict protection. In general, however, patterns of butterfly diversity are rather congruent with patterns for vascular plants and other key organism groups (Brunbjerg et al. 2018; Westgate et al. 2017).

We did not find evidence for a beneficial effect of legal protection on the survival probability of butterfly populations, neither for the period 1993-2015, when Clause 3 of the Danish Nature Protection Act was newly instated and the Natura 2000 areas being rolled out, nor for the period 2015-2019, when the said legal actions should have had sufficient time to halt biodiversity loss. This chilling observation suggests that the overall efficiency of the network of protected sites is low, something that has been found repeatedly in neighbouring countries (Kindvall et al. 2022; Rada et al. 2019). The present analysis was not designed to pinpoint the causes for this failure, but likely explanations fall in two groups: 1) too small and too isolated areas being designated for protection and 2) misguided habitat management regimes, although we cannot in principle rule out regional drivers affecting all sites in similar ways. Extinction of butterfly populations from sites, in which apparently suitable habitat has been maintained throughout the study period, could be explained by increased isolation of sites, due to destruction or deterioration of sites not included in the network of protected sites. The consequence would be lower recolonization rates, rather than increased local extinction rates at the focal sites compared to a pre-1993 benchmark. This would be aligned with the population structure of many of the studied species, which form meta-populations where species’ regional survival depends on frequent recolonization (Hanski et al. 1994), in particular for intermediately mobile species (Thomas 2000).

Habitat quality is of pivotal importance to maintain butterfly populations (Habel et al. 2022a). In Denmark, protected nature areas generally suffer from either encroachment by scrub and tall herbs due to grazing abandonment or from intense summer grazing, which depletes floral resources for anthophilous insects such as butterflies (Ejrnæs et al. in prep). This overall situation is driven by agricultural economic rationale and further incentivized by the EU common agricultural policy (Kindvall et al. 2022). Conservation areas grazed year-round at near-natural herbivore densities have been shown to better support species-rich butterfly communities (Konvička et al. 2021). One effect of over-grazing and cutting by machinery is homogenization in terms of vegetation structure and microtopography, which may increase the probability of local extinction in years with weather anomalies. Thus, the two groups of explanations outlined above are not completely separate. In any case, we deem the combined effect of them the likely cause for the demonstrated inefficacy of protected areas in terms of sustaining viable butterfly populations. It may be noted that seven species (*aquilonaris, arion, idas, optilete, rubi, semiargus and tages*) by 2019 were present only at sites under protection by Clause 3 of the Danish Nature Protection Act. This, however, was clearly an effect of initial selection of valuable sites for legal protection, rather than a sign of efficiency of the PA network.

Our study underlines the importance of a network of protected areas harbouring near-intact ecosystems at an appreciable share of total land area and being reasonably well-connected, in order to achieve sustainable conservation of butterflies, and probably other organisms as well.

## Notes

https://doi.org/10.17894/ucph.0e4f9d38-f107-42a2-b89b-5875ee05fed1

### Competing Interest Statement

The authors have declared no competing interest.

